# MRN-ATM Pathway Activation in CD4 T-Cell Senescence during Chronic Hepatitis B Virus Infection

**DOI:** 10.64898/2026.03.15.711849

**Authors:** Xiaoying Deng, Xiaoyan Wang, Yi Li, Fangfang Li, Jun Xiong, Hongyan Shi, Yun Zhou, Chuantao Ye, Xuyang Zheng, Jianqi Lian, Chao Fan, Ying Zhang

## Abstract

T-cell senescence is a hallmark of immune dysfunction in persistent viral infections, characterized by DNA damage accumulation and telomere erosion. However, the mechanisms driving CD4 T-cell senescence in the context of chronic hepatitis B virus (HBV) infection remain poorly defined. In this study, we demonstrated that people with chronic HBV infection exhibited CD4 T-cell senescence, marked by elevated KLRG1, along with increased DNA damage and telomere shortening, compared to HS. Notably, activation of the MRN-ATM (MRE11/RAD50/NBS1-Ataxia Telangiectasia Mutated Protein) pathway was prominent in CD4 T cells from HBV patients. Importantly, suppression of MRN attenuated ATM phosphorylation and its downstream signaling molecules, and inhibition of ATM reduced the production of proinflammatory cytokines in CD4 T cells derived from both HBV patients and HS. These results suggest that in chronic HBV infection, the virus induced CD4 T-cell senescence, telomere erosion, and DNA damage, while concurrent activation of the MRN-ATM pathway may serve as a compensatory mechanism to preserve CD4 T-cell function. Elucidating this relationship between T-cell senescence and DNA damage repair helps to understanding the mechanisms underlying HBV persistence and providing potential therapeutic targets against chronic HBV infection.

## Introduction

Hepatitis B virus (HBV) infection poses a significant global public health challenge, affecting more than 292 million people living with chronic hepatitis B (CHB)^[1]^. Despite available therapeutic treatments, HBV-related complications including hepatic fibrosis, liver cirrhosis, and hepatocellular carcinoma continuously contribute to substantial annual mortality^[1]^. The mechanisms of persistent HBV infection remain incompletely understood, presenting significant obstacles to achieving sustained CHB resolution.

CD4 T cells serve as central coordinators of adaptive immunity against HBV through their dual capacity to activate HBV-specific B lymphocytes and CD8 cytotoxic T cells. Impairments in adaptive immune activation and functional exhaustion of virus-specific lymphocytes constitute key determinants of persistent HBV infection^[2]^. With aging, the naive T-cell subset shrinks along with the accumulation of highly differentiated dysregulated memory cells, which makes older adults more susceptible to infections^[3]^. Recent findings suggest the premature aging in naive CD4 T-cells occur in individuals with persistent infections of human immunodeficiency virus (HIV) and hepatitis C virus (HCV)^[4–5]^. However, research on the CD4 T-cell senescence in individuals with CHB remains poorly reported.

Telomere shortening represents a cardinal feature of cellular aging. Telomeres are repetitive nucleotide-sequences of nucleotides that serve to safeguard the terminal regions of chromosomes from degradation or fusion with neighboring chromosomes^[6]^. Rapid rate of cell proliferation out of regulatory capacity of the enzyme telomerase results in telomere shortening^[6]^. A recent investigation revealed that the telomere erosion caused by telomerase deficiency led to various alterations in CD4 T cells seen in aged human subjects, including variations in naive T-cell population, CD28 expression, and cytokines production^[7]^.

Double-stranded breaks (DSBs), along with the subsequent activation of the DNA damage response (DDR), serve as fundamental drivers for the onset of cellular senescence^[8]^. The DDR cascade comprises three phases: damage sensing, signaling initiation, and repair execution^[8]^. The MRN complex, which consists of meiotic recombination 11 homolog A (MRE11A), RAD50, and Nijmegen breakage syndrome 1 (NBS1), acts as the principal DSB sensor^[9]^. Microscopically visible DNA damage foci formation, marked by local accumulations or modifications of DDR proteins (including γH2AX, 53BP1 et al), signifies repair initiation^[10]^. DNA repair amplification involves activation of ataxia telangiectasia mutated (ATM) kinase, a phosphoinositide 3-kinase (PI3K)-related kinase family member and master regulator of DNA repair signaling^[11]^. Phosphorylation of downstream effectors including CHK1, CHK2, p53 and BRCA1 mediated by activated ATM coordinates the DNA damage repair processes^[12]^.

In this study, we investigated the senescence markers and the aging phenotype of CD4 T cells in people with CHB versus HS. Then, we focused on DNA damage and activation of the MRN-ATM pathway, and demonstrated the importance of the MRN-ATM pathway in CD4 T-cell functional maintenance. Additionally, we reconfirmed the role of the MRN complex (especially NBS1) in ATM activation.

## Results

### Senescent markers were significantly higher in CD4 T cells from people with CHB compares to HS

To investigate HBV-induced senescence of CD4 T cells, we analyzed frequencies of total, naive, and memory CD4 T-cell subsets (Fig. 1a), as well as the expression of senescence markers, in individuals with CHB compared to age-matched healthy subjects (HS). People with CHB showed a trend toward decreased memory CD4 T-cell frequency (*P* = 0.079) (Fig. 1b-d), despite the fact that total and naive CD4 T-cell counts remained comparable between the two groups. T-cell aging typically manifests through decreased co-stimulatory molecules (CD27/CD28) and elevated senescence markers (CD57/KLRG1)^[13]^, along with additional findings of elevated CD38 and HLA-DR expression^[14–15]^. FCM analysis revealed a substantial increase in KLRG1 (*P*=0.0013) and a marginal rise in HLA-DR (*P*=0.068) expression in CD4 T cells derived from CHB patients (Fig. 1e-j). However, the levels of CD27, CD28, CD57 and CD38 remained the same between the two groups. Notably, the frequency (%) of KLRG1^+^ CD4 T cells showed a positive correlation with serum levels of alanine aminotransferase (ALT) in CHB patients (Fig. 1k). No significant correlations were observed between KLRG1 and other CHB markers, such as aspartate transaminase (AST), HBV DNA, HBsAg, or HBeAg (data not shown).

**Fig. 1.**
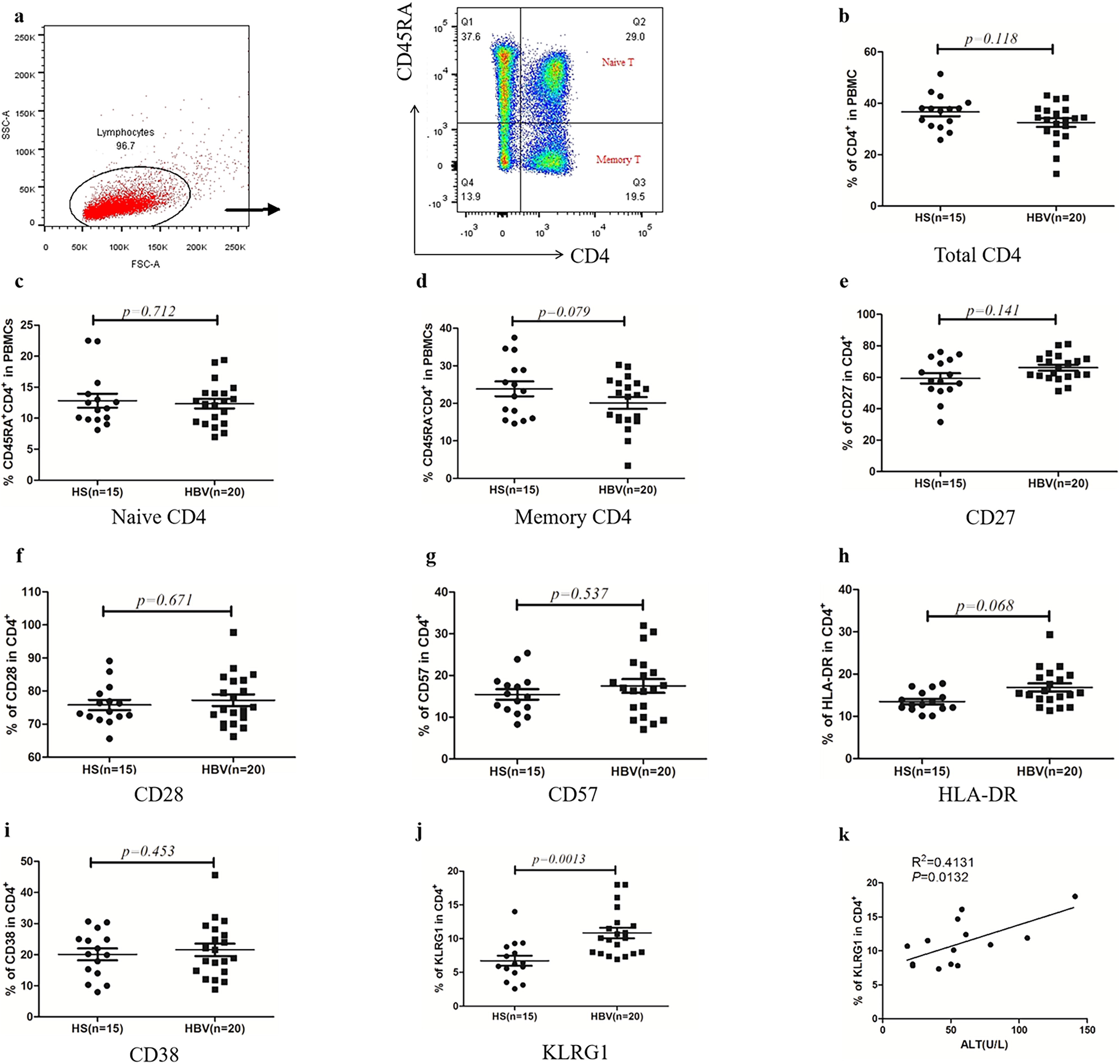
Senescent markers were elevated in CD4 T cells from people with chronic HBV infection. **a** CD4 T-cell gating strategy and dot plots of cell clustering. Representative dot plots of the FCM for the percentages of total CD4+, CD45RA+CD4+ (naive), and CD45RA−CD4+ (memory) T-cell frequencies in PBMCs are shown. **b-d** Summary data of the FCM for the percentages of total, naive and memory CD4 T-cell frequencies in PBMCs. **e-j** Summary data of the FCM for the percentages of CD27, CD28, CD57, HLA-DR, CD38 and KLRG1 levels in CD4 T cells. **k** Spearman’s correlation analysis between KLRG1 expression and ALT levels in CD4 T cells from patients. n=number of subjects studied in each group. *P* value with significant changes are shown.

### Telomere attrition and DNA damage were prominent in CD4 T cells from people with CHB compared to HS

To determine the significance of telomere attrition in cellular senescence^[16]^, we conducted a further investigation into CD4 T-cell aging dynamics by measuring telomere length in CD4 T cells isolated from CHB patients and age-matched HS using flow-FISH (Fig. 2a-d). We found that patients exhibited significantly shorter telomeres in CD4 T cells compared to HS (Fig. 2a-b; *P*<0.001). Remarkably, this telomeric shortening was consistently observed in both naive and memory CD4 T-cell subsets (Fig. 2c-d; *P*<0.001). To assess DNA damage as a contributor to T-cell senescence, γ-H2AX level in CD4 T cells was quantified through FCM and western blotting (WB). A significantly elevated γ-H2AX level was found in CHB patients compared to HS (Fig. 2e-f; *P*<0.05). To further validate DNA damage in CD4 T cells from patients, we examined colocalization of 53BP1 and r-H2AX in CD4 T cells from CHB patients versus HS by confocal microscope and found that patients demonstrated a significant increase in nuclear foci compared to HS (Fig. 2g). These results suggest prominent DNA damage and accelerated telomere erosion in CD4 T cells from patients with CHB.

**Fig. 2.**
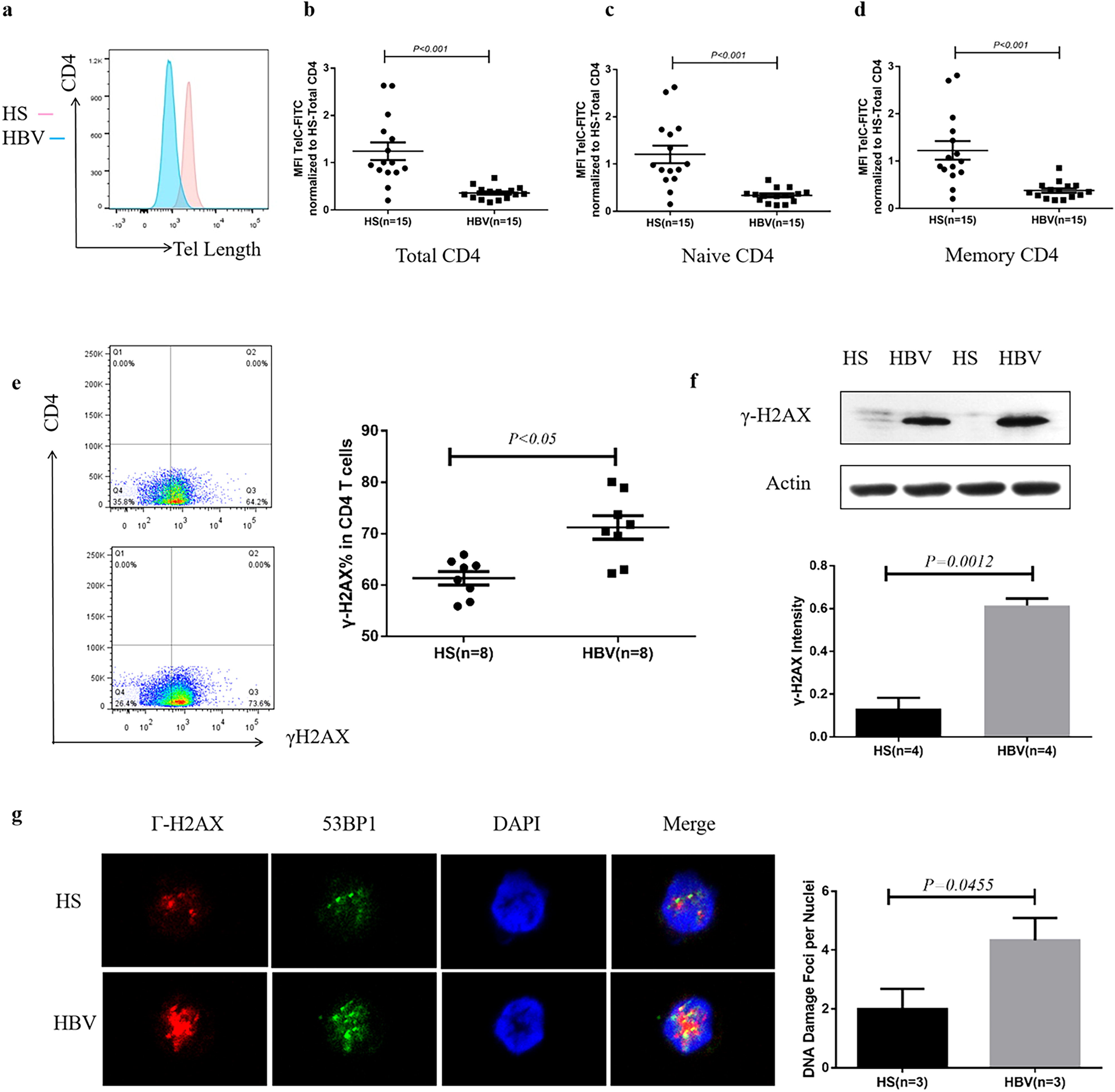
Telomere attrition and DNA damage were elevated in CD4 T cells from people with chronic HBV infection. **a** Telomere length is characterized by MFI of TelC-FITC. **b-d** Summary data of the FCM for telomeric length of total, naive and memory CD4 T-cell frequencies in PBMCs from patients and HS. **e** Representative image and summary data of the FCM for the percentages of γ-H2AX level in CD4 T cells. **f** Representative image and summary data of WB analysis for the level of γ-H2AX in CD4 T cells. **g** Representative confocal microscopic images and summary data of co-localization of 53BP1 and γ-H2AX in nuclei (DNA damage foci).

### The MRN/ATM pathway was significantly activated in CD4 T cells from people with CHB

Telomeric erosion and DNA damage were observed in CD4 T cells from CHB patients, so we further explored the activation of the DNA damage repair pathway^[17–19]^. First, we measured MRE11, RAD50, and NBS1 protein levels in CD4 cells from patients and HS. No difference was detected in the levels of MRE11 and RAD50 in CD4 T cells between two the groups (Fig3. a). However, an elevated NSB1 expression in patients was found (*P*=0.001), suggesting partial activation of the DNA damage sensors. Notably, FCM and WB analyses revealed substantially higher levels of phosphorylated ATM (p-ATM) in CD4 T cells from patients versus HS (*P*=0.0001 and *P*=0.0104, respectively; Fig. 3b-c); whereas there was no significant difference in total ATM levels between the two groups (Fig. 3c). Consistent with ATM activation, its downstream proteins, including p53, p-p53, p-CHK2, exhibited marked elevation in CHB patients (*P*=0.0396, *P*=0.0034, *P*=0.0101, respectively; Fig. 3d-f), whereas CHK2, BRCA1, and p-BRCA1 protein levels remained unchanged (Fig. 3f-g). These results indicate activation of the MRN-ATM pathway and its downstream signaling molecules.

**Fig. 3.**
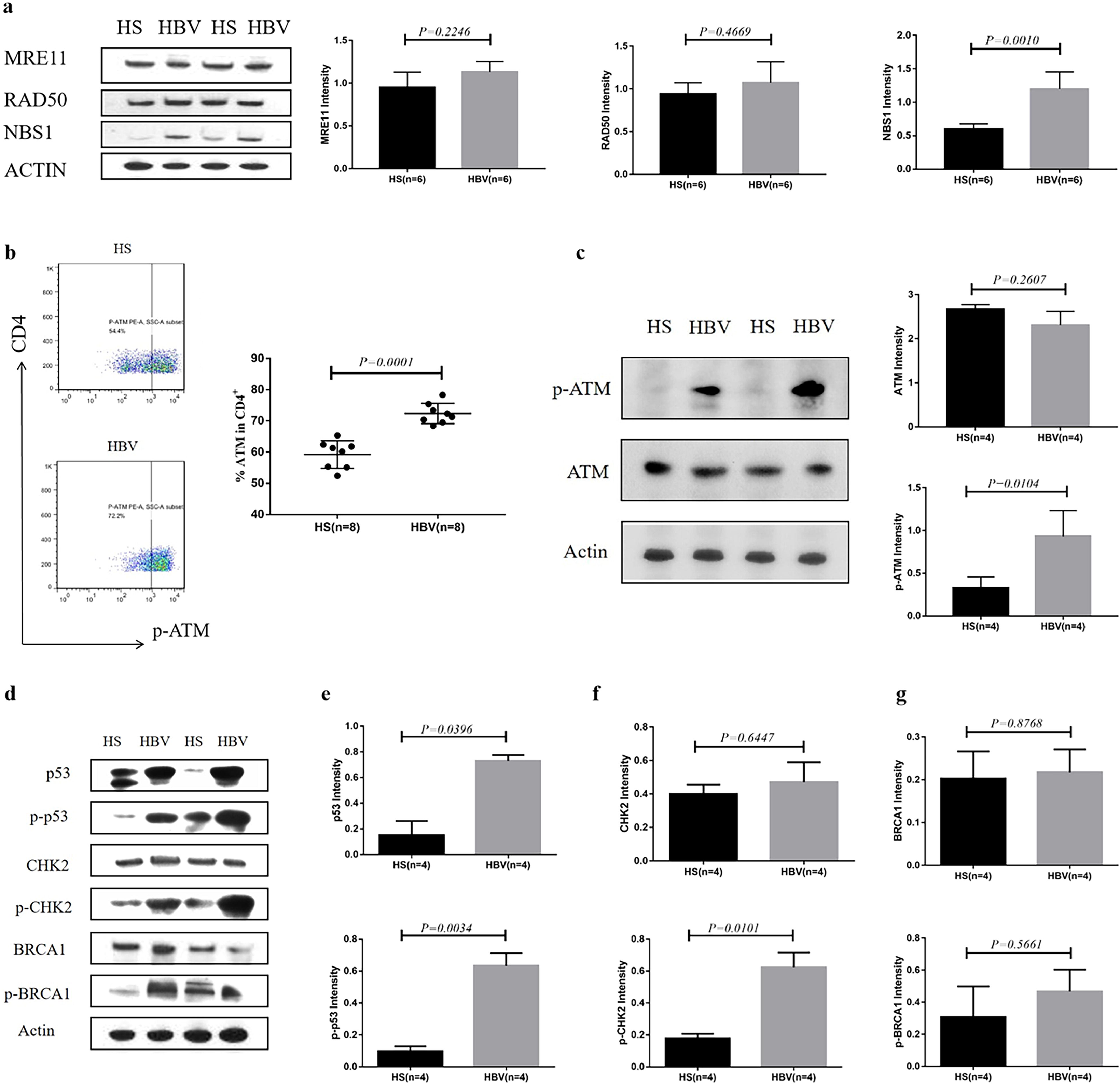
The ATM pathway was obviously activated in CD4 T cells from people with chronic HBV infection. **a** Representative image and summary data of WB analysis for the levels of MRE11, RAD50 and NBS1 in CD4 T cells isolated from patients and HS. **b** Representative dot plots and summary data of the FCM for the percentages of the level of p-ATM in CD4 T cells from patients and HS. **c** Representative image and summary data of WB analysis for the level of ATM and p-ATM. **d-g** Representative image and summary data of WB analysis for the levels of p53, p-p53, CHK2, p-CHK2, BRCA1 and p-BRCA1.

### Inhibition of ATM phosphorylation contributed to CD4 T-cell dysfunction from patients with CHB

Emerging evidence indicates that T-cell phenotype alterations during chronic viral infection are characterized by progressive T-cell dysfunction^[20]^. The ATM signaling pathway plays a critical role in repairing DNA damage, thereby mitigating cellular senescence to preserve T-cell functions^[21]^. To examine the role of ATM pathway in sustaining CD4 T-cell functions during chronic HBV infection, isolated CD4 T cells from patients were treated with a specific ATM inhibitor (KU60019), followed by measuring the proinflammatory cytokines released by CD4 T cells. We demonstrated that CD4 T cells exposed to KU60019 exhibited significant reductions in p-ATM (*P*=0.0027) and p-CHK2 (*P*=0.0002) compared to DMSO controls (Fig. 4a-c). Because IL-2 and IFN-γ are typical type-I inflammatory cytokines for assessing T-cell functions^[22]^. we analyzed their levels by ELISPOT and demonstrated that the amounts of IL-2 and IFN-γ secreted by CD4 T cells exposed to KU60019 were significantly decreased in both CHB patients and HS (*P*<0.0001; Fig. 4d-e).

**Fig. 4.**
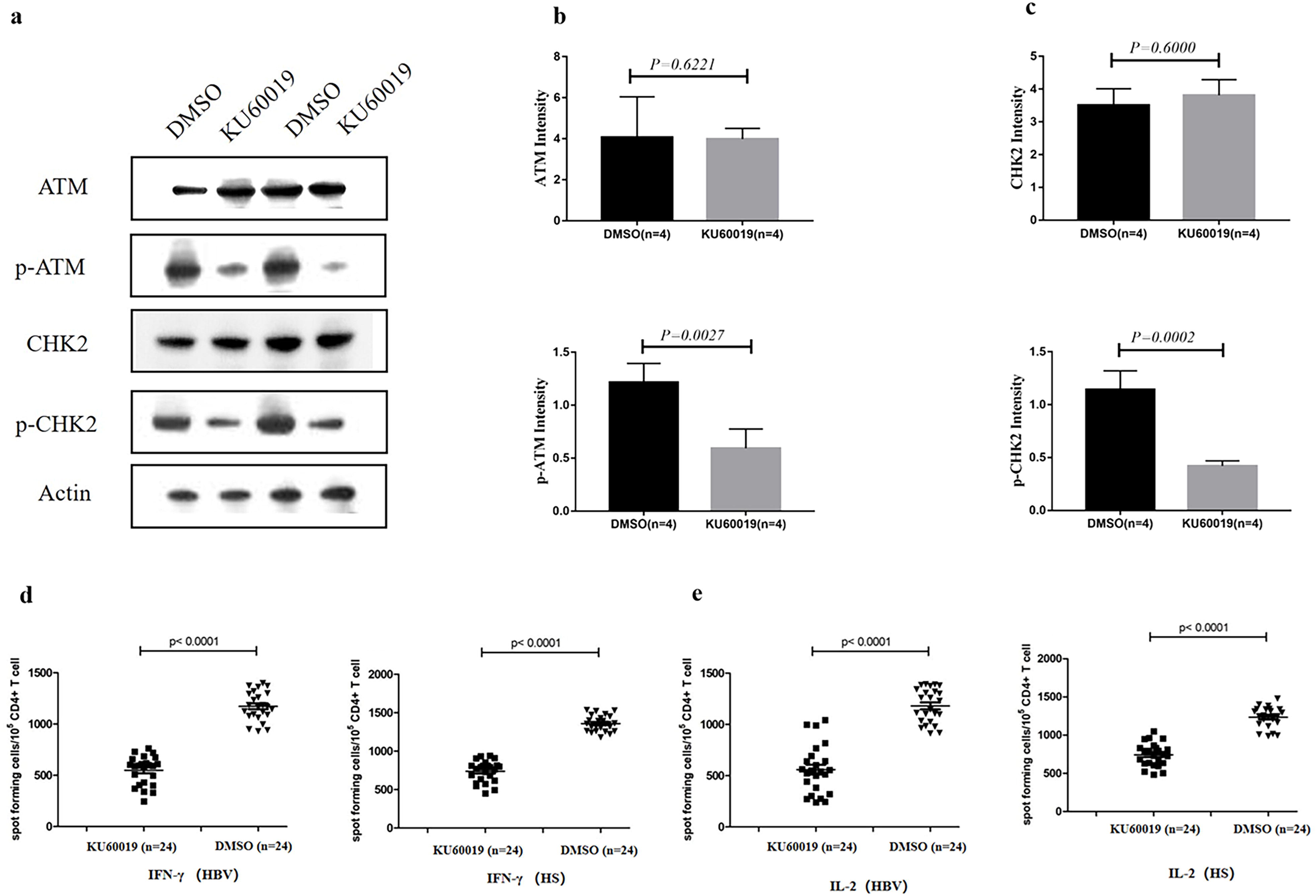
KU60019 induced CD4 T-cell dysfunction via inhibiting ATM pathway in people with chronic HBV infection and HS. a-c. Representative image and summary data of WB analysis for the levels of ATM, p-ATM, CHK2 and p-CHK2 in CD4 T cells isolated from patients which were exposed to KU60019 in advance and DMSO control. **d-e** ELISPOT test for the levels of IFN-γ and IL-2 in CD4 T cells from patients and HS with different treatments.

### Suppression of MRN complex inhibited ATM pathway in CD4 T cells from people with CHB

Given that NBS1 level was significantly elevated in CD4 T cells from CHB patients (Fig. 3a), we next determined whether blocking NBS1 may interfere with ATM activation and thus serve as a potential therapeutic target. Mirin serves as a specific inhibitor of the MRN complex^[23–24]^ and was used to inhibit NBS1 activity. We found that Mirin inhibited the expression of NBS1 (*P*=0.0033; Fig. 5a) in a dose-response manner (Fig. 5b). To exclude the possibility of non-specific effect of the Mirin in NBS1 inhibition, we used siNBS1 as a parallel control (Fig. 5c). Subsequent analysis revealed marked reduction in p-ATM and its downstream effector p-CHK2 in NBS1-inhibited/silenced CD4 T cells from patients versus control groups (Fig.5d-f); whereas there was no significant difference in the levels of total ATM and CHK2 proteins between the experimental groups and the control groups (Fig. 5e-f).

**Fig. 5.**
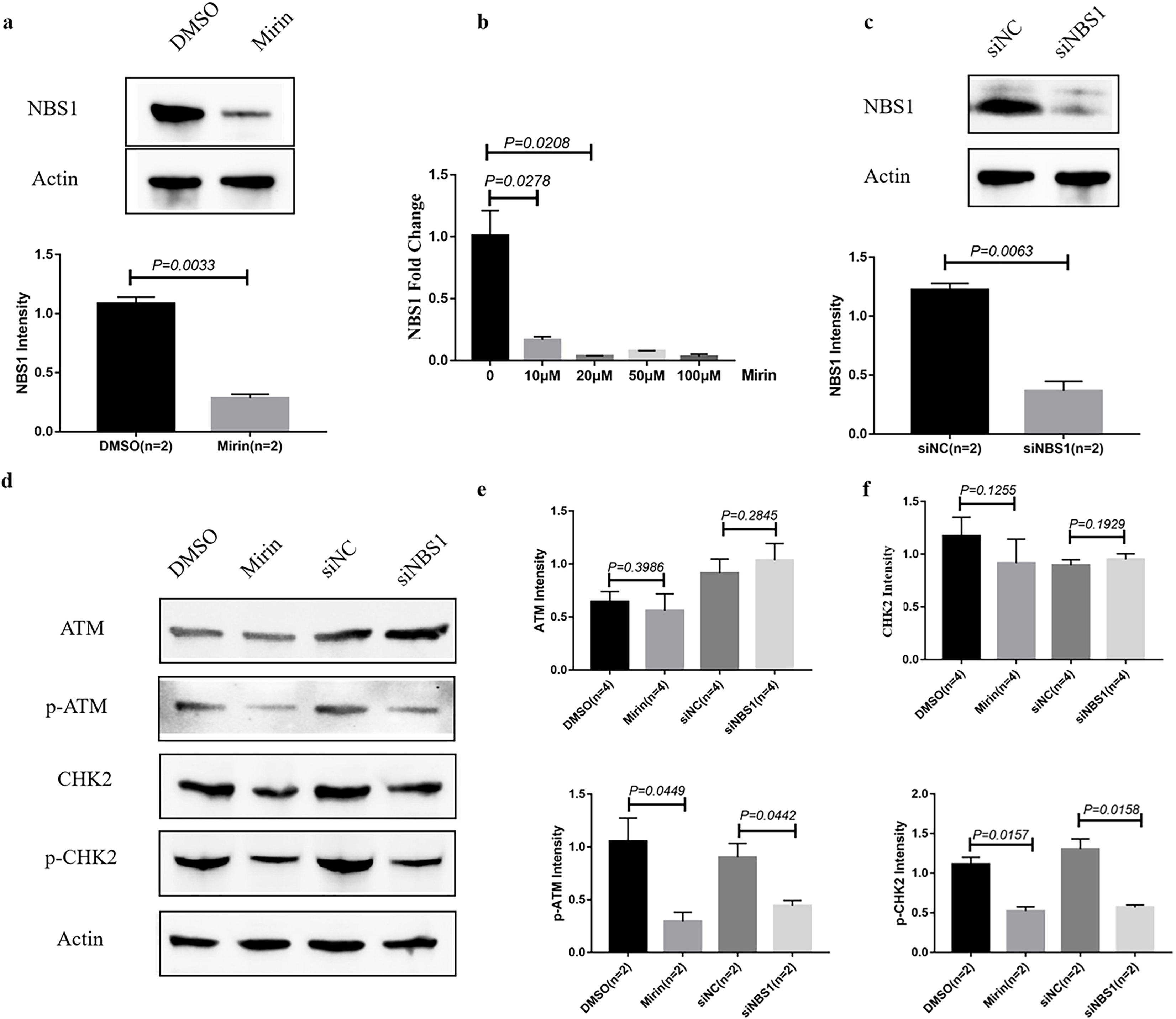
Suppression of MRN complex blocked ATM pathway in CD4 T cells from those with chronic HBV infection. **a** Representative image and summary data of WB analysis for the levels of NBS1 in CD4 T cells purified from patients which were exposed to Mirin in advance and DMSO control. **b** Investigation of optimal concentration of Mirin (Optimal concentration: 10μM, P=0.0278). **c** Representative image and summary data of WB analysis for the levels of NBS1 in CD4 T cells isolated from patients with the treatment of siNBS1 and siNC control. **d-f** Representative image and summary data of WB analysis for the levels of ATM, p-ATM, CHK2 and p-CHK2 in CD4 T cells isolated from patients with different treatments.

## Discussion

In this study, we examined the senescence markers of CD4 T cells in people with CHB versus HS and demonstrated an aging phenotype, including elevated KLRG1 level and shortened telomere length, in CD4 T cells in patients with CHB. Additionally, we found a prominent DNA damage and activation of the MRN-ATM pathway. Importantly, inhibition of ATM activation reduced IL-2 and IFN-γ secretion in CD4 T cells, indicating the importance of the ATM pathway in CD4 T-cell functional maintenance. Also, suppression of the MRN/NBS1 resulted in a decrease in p-ATM and its downstream molecules, reconfirming an essential role of the MRN complex (especially NBS1) in ATM activation (Fig. 6) and possible target for therapeutic intervention.

**Fig. 6.**
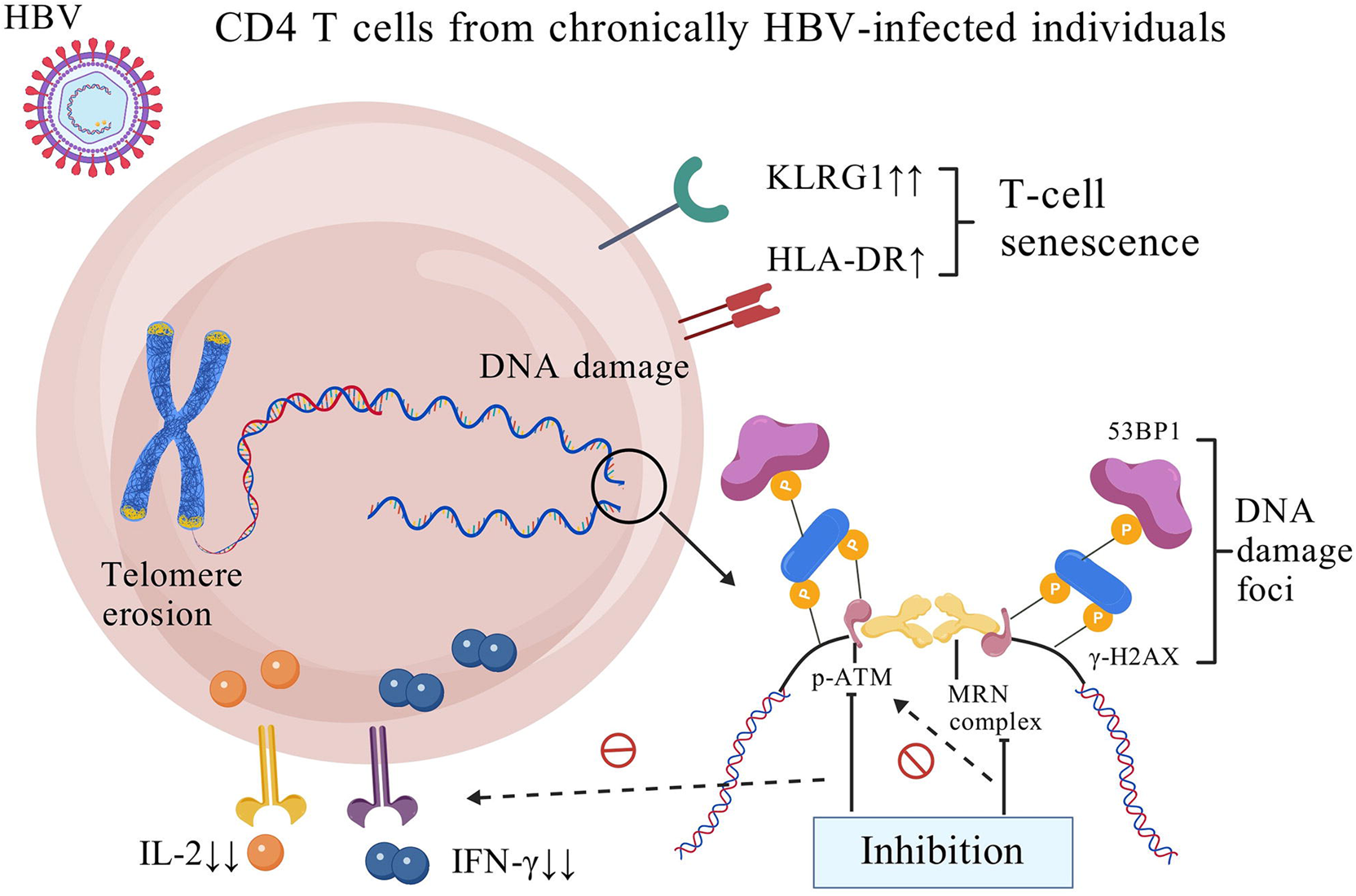
**Schematic model.**

T-cell senescence has been extensively documented in HIV and HCV infections, characterized by upregulation of senescence markers^[4,5,25]^. Our observation of KLRG1 elevation – without significant alterations in canonical markers (CD27, CD28, CD57, CD38)^[26–27]^ – highlighted the distinctive characteristics of HBV infection in T cell senescence. Notably, KLRG1 correlated with serum ALT levels in CD4 T cells from CHB patients, suggesting a potential link between HBV-induced T-cell senescence and hepatic injury. Several studies demonstrated that senolytic agents might synergize with antiviral therapies to restore immune competence^[28]^. Thus, our findings may carry clinical relevance for chronic HBV infection.

Cellular senescence can be triggered by unrepaired DNA damage that forces cells into growth arrest and/or death^[29]^. Unlike the decrease in ATM activation observed in other chronic viral infections^[4–5]^, the pronounced ATM activation was observed in CD4 T cells from people with CHB. Based on clinical parameters, we noted that many CHB patients in our cohort maintain relatively normal hepatic function and morphology despite persistent HBV infection. This may involve protective effects of the ATM pathway on immune cells. Further analysis of the ATM status in CHB patients with different clinical characteristics (stratified subgroups) are needed to fully understand this discrepancy. Our findings with inhibition of the ATM pathway resulted in decreased cytokine secretion by CD4 T cells also demonstrated the protective role of ATM in T-cell functions. Although ATM pathway mitigated functional decline by facilitating DSB repair, its inability to fully resolve genomic instability suggested its intrinsic limitations in reparing capacity. Additionally, some studies suggested that ATM activation regulated senescence via lysosomal-mitochondrial axis regulation and autophagy induction^[30]^. Our team is currently investigating the relationships between mitochondrial autophagy and ATM activation in CD4 T cells from people with CHB.

MRN complex provides novel perspectives on therapeutic strategies for its essential role in activation of the ATM pathway. A recent study has discovered that knockdown of MRN complex significantly inhibited the formation of cccDNA in HBV infection^[31]^, suggesting the potential of MRN inhibitors as therapeutic drugs for chronic HBV treatment. Additionally, the replication of minute virus of mice (MVM) depends on MRE11^[32]^; RAD50 expression is correlated with cancer progression and poor survival in people with HBV-related HCC^[33]^. Therefore, MRN complex should be paid more attention on the potential of antiviral drug development.

Despite extensive efforts in this study, certain limitations and unresolved issues remained. First, *in vitro* CD4 T-cell models may not fully recapitulate the *in vivo* microenvironment, particularly interactions with hepatocytes or regulatory T cells within liver where HBV resides. What’s more, whether HBV directly induces DNA damage via viral proteins or indirectly through chronic inflammation requires more exploration.

In conclusion, we believe that the MRN-ATM axis may establish a balance between viral persistence and T-cell antiviral immunity, providing a conceptual framework for elucidating the role and mechanisms of ATM pathway in T cell senescence and HBV persistence as well as potential therapeutic targets.

## Materials and methods

### Subjects

A total of 84 participants were enrolled in this study, of whom 45 had chronic HBV infection and 39 were healthy age-matched. Information on all participants and sample usages are listed in Table S1. Chronic HBV infection was defined as the presence of HBsAg in serum for over 6 months and individuals with liver cirrhosis were excluded. None of the patients received antiviral therapy. All individuals were negative for antibodies to hepatitis A virus (HAV), HCV, hepatitis D virus (HDV), and HIV. All subjects included in the study were sourced from Department of Infectious Diseases, Tangdu Hospital, The Fourth Military Medical University (Xi’an, China).

### Cell isolation and culture

PBMCs were isolated from whole blood by Lymphoprep^TM^ (STEMCELL, Bernburg, Germany) density centrifugation. CD4 T cells were isolated from PBMCs using CD4^+^T Cell Isolation Kit and a MACS Separator with LS Column (Miltenyi Biotec, Cologne, Germany). The isolated T cells were cultured in RPMI 1640 medium (Pricella, Wuhan, China) at 37℃ and 5% CO2 atmosphere.

### Flow cytometry

In the analysis of CD4 T-cell subsets, PBMCs were stained with CD4-APC, CD45RA-FITC (BD, Franklin Lakes, NJ, USA) antibodies, or IgG1 k Isotype Control and IgG2b k Isotype Control (BD). For intracellular staining, the cells were fixed and permeabilized with BD Cytofix/Cytoperm™ Fixation/Permeabilization Kit (BD), and stained with Anti-Human CD27, Anti-Human CD28, Anti-Human CD57 Anti-Human CD38 Anti-Human HLA-DR, Anti-Human KLRG1 (BD), and ATM-FITC antibody (Abcam, Cambridge, UK), p-ATM-FITC antibody (Abcam), and γ-H2AX antibody (Thermo Fisher Scientific/TMO, Waltham, MA, USA). The stained cells were analyzed on Accuri^TM^ C6 flow cytometer (BD), and data were analyzed by FlowJo 7.6 software (BD). Isotype control antibodies (eBioscience) and fluorescence minus one controls were used to determine the background levels of staining and adjust multicolor compensation as gating strategy.

### Flow-FISH

The Flow-FISH (fluorescence in-situ hybridization) method was used to measure telomere lengths. T cells were subjected to incubation in a hybridization buffer with 0.25 μM of FITC-PNA Tel C probe (CCCTAAC repeats) (PNA Bio, Thousand Oaks, CA, USA) for 10 min at room temperature. The samples were heated for 10 min at 85°C, chilled on ice, and hybridized in the dark at room temperature overnight. The samples were analyzed by FCM, and CD4 T-cell telomere lengths were reported as mean fluorescence intensity (MFI).

### Western blotting

CD4 T cells were lysed on ice using radio immunoprecipitation assay (RIPA) lysis buffer (Beyotime Biotechnology, Shanghai, China) in the presence of phosphorylated protease inhibitor and phenylmethanesulfonyl fluoride (PMSF) (100mM) (Beyotime). The protein concentrations were determined using Pierce BCA protein assay kit (Beyotime). Proteins were separated by SDS-PAGE (Beyotime), transferred to polyvinylidene difluoride (PVDF) membranes (TMO), which were blocked with 5% nonfat milk, 0.5% Tween-20 (TMO) in TBS (Beyotime), and incubated with the ATM (D2E2), pATM (Ser1981) (D6H9), CHK2 (D9C6), p-CHK2 (Thr68) (C13C1), p53, p-p53 (Ser15), BRCA1, p-BRCA1, Rad50 (E3I8K), NBS1, β-Actin antibodies (Cell Signaling Technology/CST, Danvers, MA, USA), and Anti-γ H2AX (phospho S139), Anti-Mre11 antibodies (Abcam). Horseradish peroxide-conjugated secondary antibodies (CST) were then used and the proteins were detected with Pierce^TM^ ECL Western Blotting Substrate (TMO).

### Confocal microscopy

Immunofluorescence staining was conducted following the established protocol. The CD4 T cells were fixed in 2% paraformaldehyde for 20 minutes, permeabilized with 0.3% Triton X-100 in PBS for 10 minutes, blocked with 5% BSA in PBS for 1 hour, and then incubated with rabbit anti-53BP1 antibody (CST) and anti-γ-H2AX (Ser-139) antibody (Biolegend, San Diego, CA, USA) at 4 °C overnight. The cells were washed with PBS with 0.1% Tween-20 for three times, and then stained with anti-rabbit IgG-Alexa Fluor 488 and anti-mouse IgG-Alexa Fluor 555 (Invitrogen, Carlsbad, CA, USA) at room temperature for 1 h, washed and mounted with DAPI Fluoromount-G (SouthernBiotech, Birmingham, AL, USA). Images were acquired with a confocal laser-scanning inverted microscope (Danaher Corporation, Washington, D.C., USA).

### Inhibition of ATM phosphorylation and MRN complex

KU-60019 (Abcam), a kind of highly selective ATM kinase inhibitor, should be soluble in DMSO (Sigma-Aldrich, Shanghai, China). Mirin (Abcam) is a MRN/ATM complex inhibitor that prevents ATM activation in response to double strand breaks in vitro. The NBS1-encoding gene was silenced by siRNA 11612 (TMO). The siNC served as the negative control reagent.

### ELISPOT

ELISPOT assays were performed using Human IFN-γ ELISPOT Kit and Human IL-2 ELISPOT kit (Abcam) with optimized protocols. Capture antibody coating: 100 μL/well of anti-IFN-γ and anti-IL-2 in 100 μL/well PBS (4°C, overnight). Spot quantification was measured by an ImmunoSpot S6 Entry M2 Analyzer (CTL, Shaker Heights, OH, USA).

### Statistical analysis

The data were expressed as mean ± standard deviation (SD) of three independent experiments. All data were analyzed using GraphPad Prism software (ver. 5.0; GraphPad Software, Inc.). One-way ANOVA or Student’s t-test was used to analyze differences between groups. A *P* value<0.05 was considered as statistically significant.

### Schematic model drawing

The schematic model was created with BioGDP.com^[34]^.

### Supplementary materials

**Table S1 Information on all participants and sample usages.**

## Declaration

### Availability of data and materials

The datasets generated and/or analysed during the current study are not publicly available due to privacy concerns but are available from the corresponding author on reasonable request.

## Supporting information

graphical abstract

## Acknowledgements

This work was supported by National Natural Science Foundation of China to Zhang Y. (81870414), by a Tangdu Hospital Funds for International Cooperation and Exchange to Zhang Y. (2017GJHZ001), and by Yizhen Genetics Talent Development Program to Deng X.Y. (2025YZ17).

## Ethics approval and consent to participate

The study protocol was approved by the institutional review board (IRB) of Tangdu Hospital (GKJ-Y-202403-105), The Fourth Military Medical University (Xi’an, China).

## Author Contributions

**Xiaoying Deng:** Conceptualization, Methodology, Formal analysis, Investigation, Data Curation, Writing - Original Draft & Editing, Visualization.

**Xiaoyan Wang:** Conceptualization, Methodology, Formal analysis, Investigation, Resources, Data Curation, Visualization.

**Yi Li:** Formal analysis, Resources.

**Fangfang Li:** Formal analysis, Resources.

**Jun Xiong:** Formal analysis, Resources.

**Hongyan Shi:** Formal analysis, Resources.

**Yun Zhou:** Formal analysis, Investigation, Resources.

**Chuantao Ye:** Formal analysis, Investigation, Resources.

**Xuyang Zheng:** Formal analysis, Investigation, Resources.

**Jianqi Lian:** Conceptualization, Validation, Project administration.

**Chao Fan:** Conceptualization, Methodology, Data Curation, Writing - Review & Editing, Project administration.

**Ying Zhang:** Conceptualization, Validation, Writing - Review & Editing, Supervision, Project administration, Funding acquisition.

