## Supplementary figures and images for "MRN-ATM Pathway Activation in CD4 T-Cell Senescence during Chronic Hepatitis B Virus Infection"

### graphical abstract

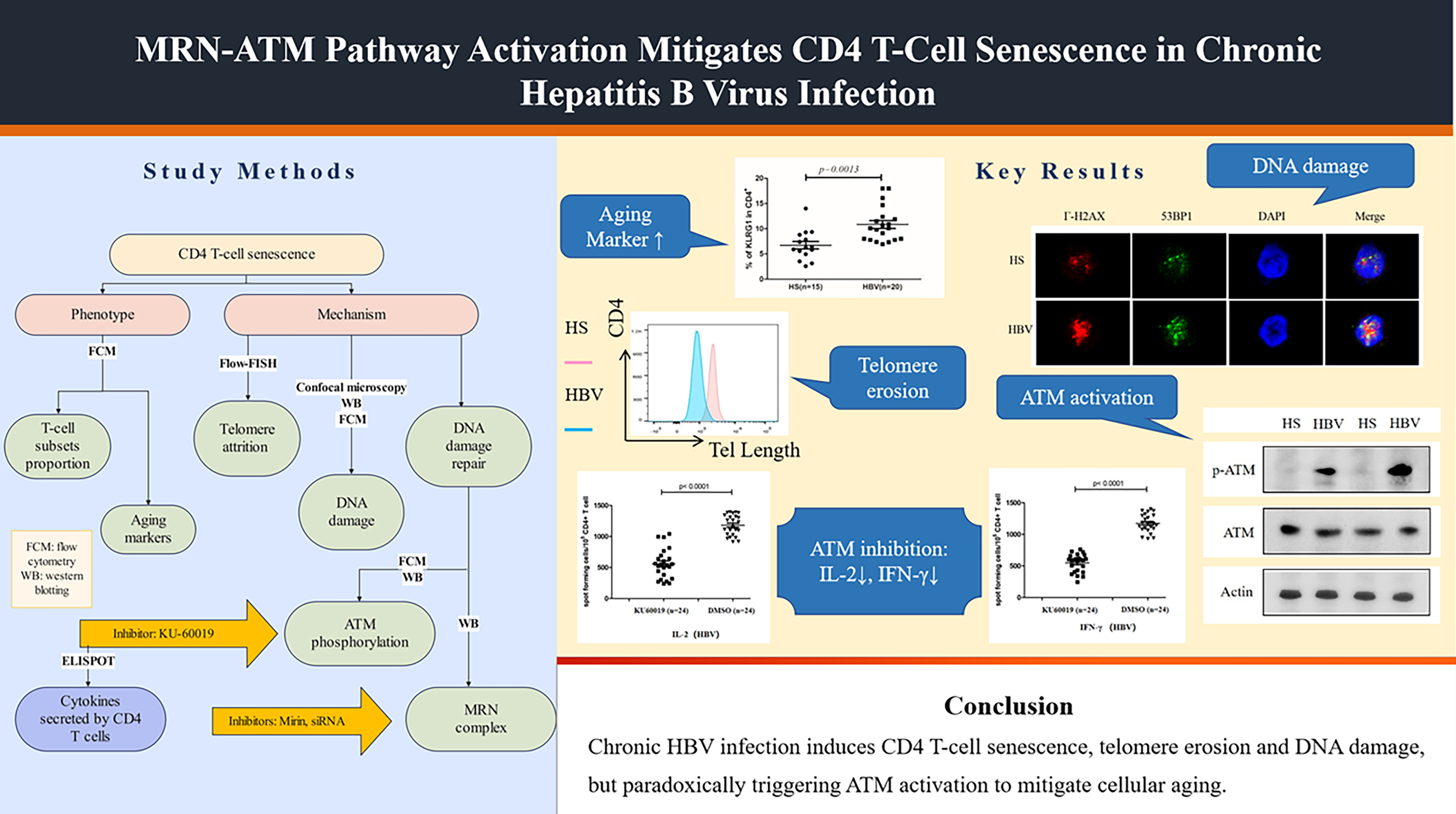
